# Mechano-modulatory synthetic niches for liver organoid derivation

**DOI:** 10.1101/810275

**Authors:** Giovanni Sorrentino, Saba Rezakhani, Ece Yildiz, Sandro Nuciforo, Markus H. Heim, Matthias P. Lutolf, Kristina Schoonjans

## Abstract

The recent demonstration that primary cells from the liver can be expanded *in vitro* as organoids holds enormous promise for regenerative medicine and disease modeling^1–5^. The use of three-dimensional (3D) cultures based on ill-defined and potentially immunogenic matrices, however, hampers the translation of liver organoid technology into real-life applications^6^. We here used chemically defined hydrogels for the efficient derivation of both mouse and human hepatic organoids. Organoid growth was found to be highly stiffness-sensitive and dependent on yes-associated protein 1 (YAP) activity. However, in contrast to intestinal organoids^7^, YAP-mediated stiffness sensitivity was independent of acto-myosin contractility, requiring instead activation of the Src family of kinases (SFKs). Aberrant matrix stiffness on the other hand led to a shift in the progenitor phenotype, resulting in compromised proliferative capacity. Finally, we demonstrate the unprecedented establishment of biopsy-derived human liver organoids without the use of animal components at any step of the process. Our approach thus opens up exciting perspectives for the establishment of protocols for liver organoid-based regenerative medicine.

## Introduction

Although the liver has a remarkable regenerative potential, chronic inflammation and scarring severely impair liver regeneration^8^, making organ transplantation the only treatment option for patients with severe liver failure^9^. This therapeutic approach, however, is limited by the lack of liver donors, emphasizing the urgent need for cell-based therapies^10^. A promising alternative to liver transplantation comes from the recent breakthrough that liver organoids can be generated *in vitro* within animal-derived matrices (e.g. Matrigel) from mouse and human bile duct-derived bi-potential facultative progenitor cells^1,2,11^ or primary hepatocytes^4,5^. These organoids are largely composed of progenitor cells that are genetically stable and can be differentiated into functional hepatocyte-like cells, which are able to engraft and increase survival when transplanted in a mouse model of liver disease^1,2^. However, the batch-to-batch variability of the three-dimensional (3D) matrices currently used for organoid derivation, as well as their mouse tumour-derived origin, makes them unsuited for therapeutic ends. Recent work has suggested that composite matrices of fibrin and laminin-111, optimized for intestinal organoid culture, could also be used for liver organoid growth^12^. Owing to the mouse-tumour derived laminin, these matrices are however incompatible with clinical use, and to the best of our knowledge there is no protocol available to expand and differentiate clinical-grade hepatic organoids^7,13^.

## Results

We previously reported chemically defined 3D matrices for intestinal stem cell culture and organoid derivation^7^, identifying design principles that could be adopted to mimic stem cell niches from different tissues. Here, we sought to develop a synthetic matrix for the efficient proliferation of liver progenitor cells by recapitulating key physical and biochemical characteristics of the hepatic microenvironment. To this aim, we first generated inert poly(ethylene glycol) (PEG) hydrogels enzymatically crosslinked by the activated transglutaminase factor XIIIa (FXIIIa)^14^. To mimic the mechanical properties of the mouse liver, we tuned the stiffness of PEG gels to physiological values (≈1.3 kPa)^15,16^. Key ECM proteins found in the native liver^17^, such as laminin-111, collagen IV and fibronectin, were then incorporated in the PEG network, and soluble factors found in the hepatic niche, such as hepatocyte growth factor (HGF)^18^, the Wnt agonist R-Spondin^2,19^ and fibroblast growth factor 10 (FGF10)^20^, were added to the culture medium, referred to as expansion medium (EM)^2^.

Single dissociated mouse liver progenitor cells derived from Matrigel-expanded liver organoids were embedded into either Matrigel or PEG hydrogels and cultured in EM (Supplementary Fig. 1a). The functionalization of PEG hydrogels with fibronectin and laminin-111 led to efficient organoid generation, comparable to Matrigel (Figs. 1a, b). Replacement of the full-length fibronectin with its minimal integrin recognition peptide RGDSPG (Arg-Gly-Asp-Ser-Pro-Gly) led to similar results, suggesting that the addition of a minimal adhesive moiety to the otherwise inert matrix is sufficient to promote extensive proliferation of liver progenitors (Figs. 1a, b). Cells expanding in PEG hydrogels modified with RGDSPG (‘PEG-RGD’) generated cystic structures characterized by a central lumen and a surrounding epithelium (Figs. 1c, e). Histology and gene expression analyses showed that these PEG-RGD-derived organoids possess a progenitor phenotype expressing stem/ductal markers such as *Lgr5, Epcam, Krt19* and *Sox9* (Figs. 1d, e) and, in terms of morphology and gene expression, are indistinguishable from organoids grown in Matrigel (Figs. 1d, e). As expected, markers of fully differentiated hepatocytes such as *Cyp3a11* were not expressed (Fig. 1d).

**Figure 1.**
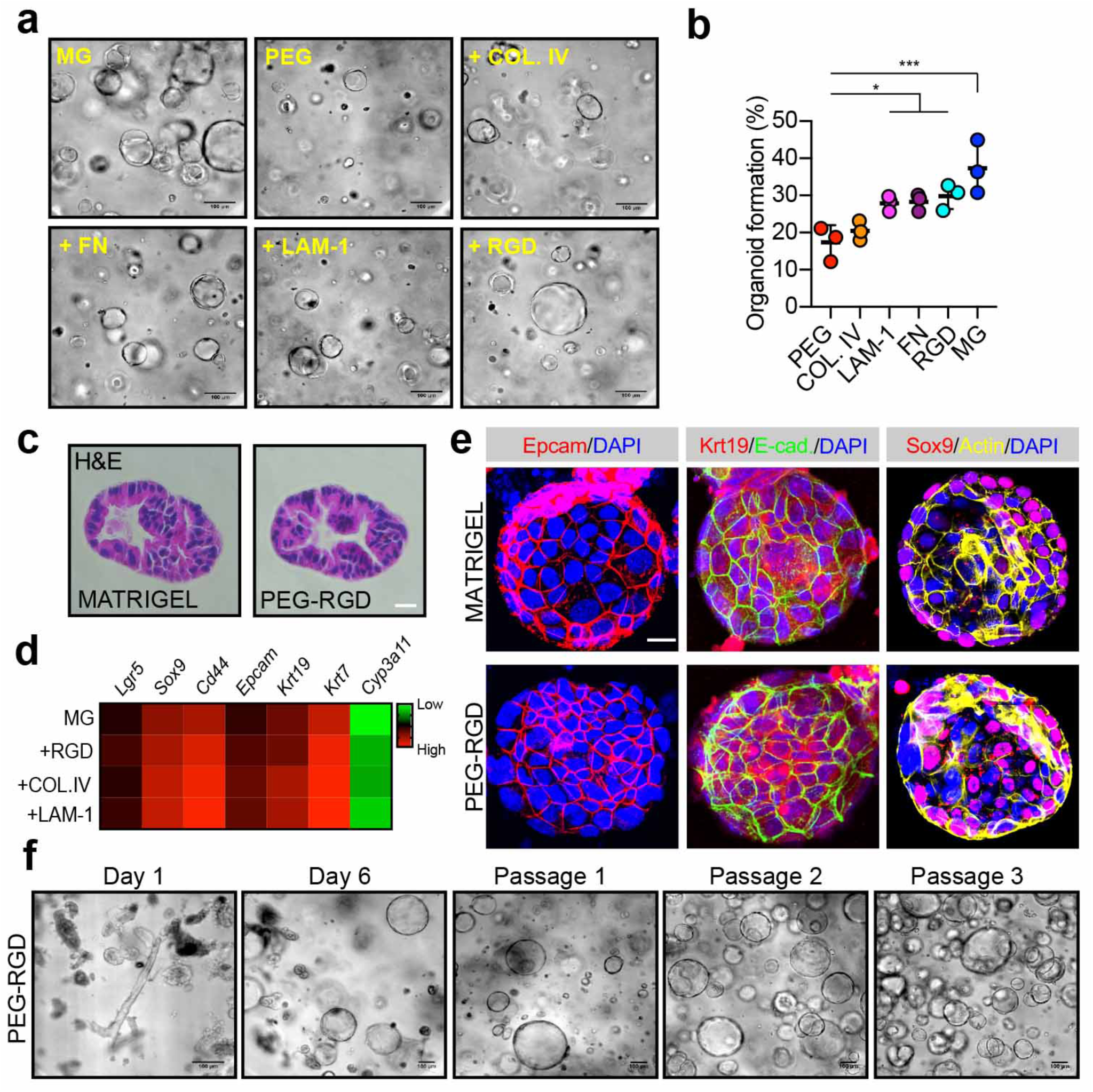
Liver organoid growth in Matrigel and PEG supplemented with ECM components. **(a)** Mouse liver progenitor cells 3 days after embedding in: Matrigel (MG), plain PEG (PEG) and PEG functionalized with the indicated ECM components: COL.IV (collagen IV), FN (fibronectin), LAM-1 (laminin-1) and RGD-representing peptide (RGD). **(b)** Quantification of organoid formation efficiency relative to panel a. **(c)** Hematoxylin and eosin staining of Matrigel- and PEG-derived organoids. Scale bar 25 μm. **(d)** Gene expression was analysed by qPCR in liver organoids 6 days after embedding in Matrigel and PEG hydrogels supplemented with different ECM factors. **(e)** Liver organoid immunostaining was performed 6 days after embedding of liver progenitor cells in Matrigel or PEG-RGD. **(f)** Liver organoids can be cultured in PEG hydrogels by directly embedding mouse biliary duct fragments. Representative pictures are shown. Graphs show individual data points derived from n=3 independent experiments and means ± s.d. *P<0.05, ***P<0.001, one way Anova.

A major limitation of all current protocols for culturing epithelial organoids is an obligatory requirement of Matrigel (or similar natural ECM-derived matrices) in the first step of organoid generation. To test whether mouse liver organoids can be established in synthetic matrices without any initial Matrigel culture step, biliary duct fragments were isolated from mouse liver and directly embedded in PEG-RGD hydrogels (Supplementary Fig. 1a). Strikingly, after 6 days of culture, organoids emerged that could be serially passaged in culture (Fig. 1f). PEG-RGD gels allowed organoid growth for more than 14 days (Supplementary Fig. 1b) without any significant structural deterioration. In contrast, Matrigel softened and no longer provided sufficient mechanical support already after 6 days of culture (Supplementary Fig.1c), highlighting the importance of having a stably crosslinked matrix for long-term organoid culture.

Liver organoids can be differentiated into functional hepatocyte-like cells when cultured in the presence of specific differentiation medium (DM) containing inhibitors of Notch and TGF-*β* pathways^2^. To test whether organoids derived in the synthetic matrix preserved the capacity for differentiation and hepatocyte maturation, we grew organoids in PEG-RGD gels and replaced the expansion medium with differentiation medium after 6 days, and analysed the expression of differentiated hepatocyte markers after 12 days^2^. Similar to Matrigel cultures, synthetic gels promoted a robust increase in the transcript levels of mature hepatocyte markers such as *Cyp3a11, Alb, Ttr, Nr1h4* (*Fxr*), *Slc2a2* (*Glut2*), *Glul* and *Nr1h3* (*Lxr*) (Fig. 2a and Supplementary Fig. 2b) and expression of ALB and HNF4*α* proteins (Fig. 2b), while markers of stem cells, such as *Lgr5*, disappeared (Supplementary Fig. 2a). As expected, during the differentiation process, cells acquired a characteristic hepatocyte-like morphology, as evidenced by a polygonal shape and expression of junction proteins such as ZO-1 and E-cadherin (Figs. 2c, d). Occasionally, hepatocyte-like cells showed poly-nucleation, a typical feature of hepatocytes (Figs. 2c, d and Supplementary Fig. 2d).

**Figure 2.**
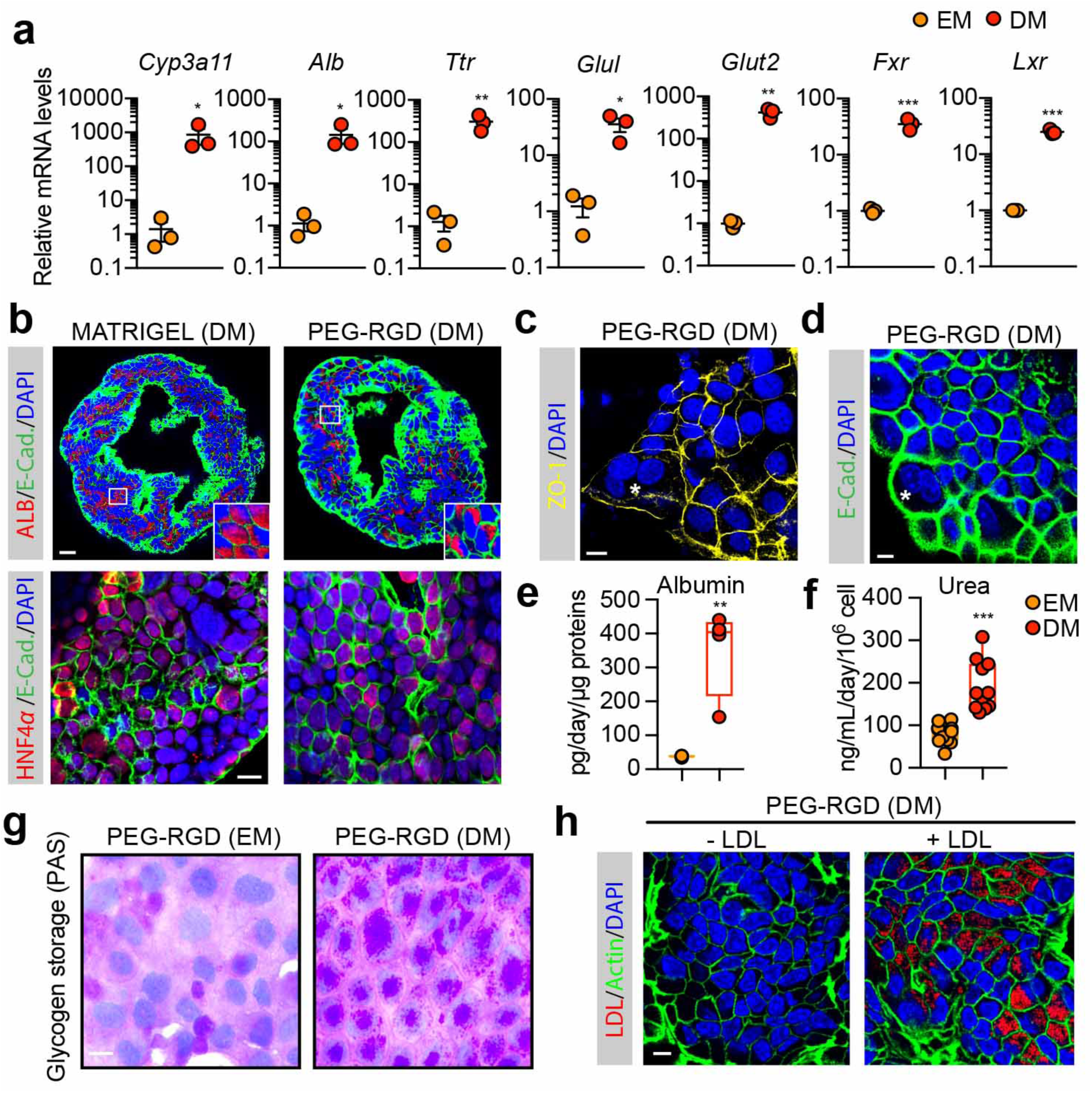
Differentiation of liver organoids into hepatocyte-like cells in PEG-RGD hydrogels. **(a)** Gene expression was analysed by qPCR in liver organoids maintained in expansion medium (EM) or differentiation medium (DM) in PEG-RGD hydrogels. Graphs show individual data points derived from n=3 independent experiments and means ± SEM. *P<0.05, ** P<0.01 ***P<0.001 paired Student’s two-tailed t-test. **(b)** Representative confocal immunofluorescence images of Albumin (ALB), Hnf4a and E-Cadherin (E-CAD.). Scale bars: 25 μm (top), 10 μm (bottom). **(c)** Representative confocal immunofluorescence images of ZO-1 in liver organoids maintained in differentiation medium (DM) in PEG-RGD hydrogels. The asterisk indicates a bi-nucleated cell. Scale bar 10 μm. **(d)** Liver organoids maintained in differentiation medium (DM) in PEG-RGD hydrogels. E-cadherin was used to visualize cell borders. The asterisk indicates a bi-nucleated cell. Scale bar 10 μm. **(e)** Albumin secretion was quantified in the supernatant of organoids embedded in PEG-RGD hydrogels and maintained in EM or DM. Box plots show min. to max. individual data points derived from n=4. **P<0.001, unpaired Student’s two-tailed t-test **(f)** Urea production was quantified in the supernatant of organoids embedded in PEG-RGD hydrogels and maintained in EM or DM. Box plots show min. to max. individual data points derived from n=11. **P<0.001, unpaired Student’s two-tailed t-test **(g)** Glycogen accumulation was assessed by PAS (Periodic-Acid Schiff) staining in liver organoids embedded in PEG-RGD hydrogels and maintained in EM or DM. Scale bar 10 μm. **(h)** LDL uptake was monitored by Dil-ac-LDL fluorescent substrate in liver organoids maintained in DM in PEG-RGD hydrogels. Scale bar 10 μm.

Next, to test whether hepatocyte-like cells generated in PEG-RGD hydrogels display hepatocyte-specific functions, we monitored albumin secretion. Liver organoids grown in DM showed a marked increase in albumin secretion, as compared to organoids cultured in EM (Fig. 2e). Moreover, hepatocyte-like cells were able to produce and secrete urea, as well as to accumulate glycogen, two enzymatically-regulated processes that occur in mature hepatocytes (Figs. 2f, g). Moreover, the majority of differentiated cells were capable of internalizing low-density lipoproteins (LDL) from the culture medium, indicating that LDL receptor-mediated cholesterol uptake is functional (Fig. 2h). Altogether, these results demonstrate that liver organoids grown in PEG-RGD hydrogels can be readily differentiated into hepatocyte-like cells that mimic many of the established hepatic functions.

Mechanical signals can play a critical role in controlling stem cell behaviour and tissue homeostasis^21^, but also contribute to the manifestation of diseases^22–24^. Despite the recent progress in establishing novel 3D liver model systems^2,3,5^, relatively little is known about the role of mechanics in regulating hepatic stem cell biology. To test whether matrix mechanics affect liver organoid growth, we grew organoids in hydrogel of variable stiffness, ranging from values below the normal mouse liver stiffness (0.3 kPa) to those reaching physiological stiffness (1.3 kPa)^15,16^. Organoid formation efficiency was profoundly affected by the mechanical properties of the matrix, with values mimicking physiological liver stiffness (between 1.3 and 1.7 kPa) being optimal (Figs. 3a, b and Supplementary Fig. 3a). Differentiation capacity however was unaffected by the degree of stiffness as induction of hepatic genes was maintained also when liver organoids were differentiated in soft gels (Supplementary Figs. 2c and 3b). These data demonstrate that optimizing the mechanical properties of the hydrogel represents a critical step in the efficient generation of liver organoids.

**Figure 3.**
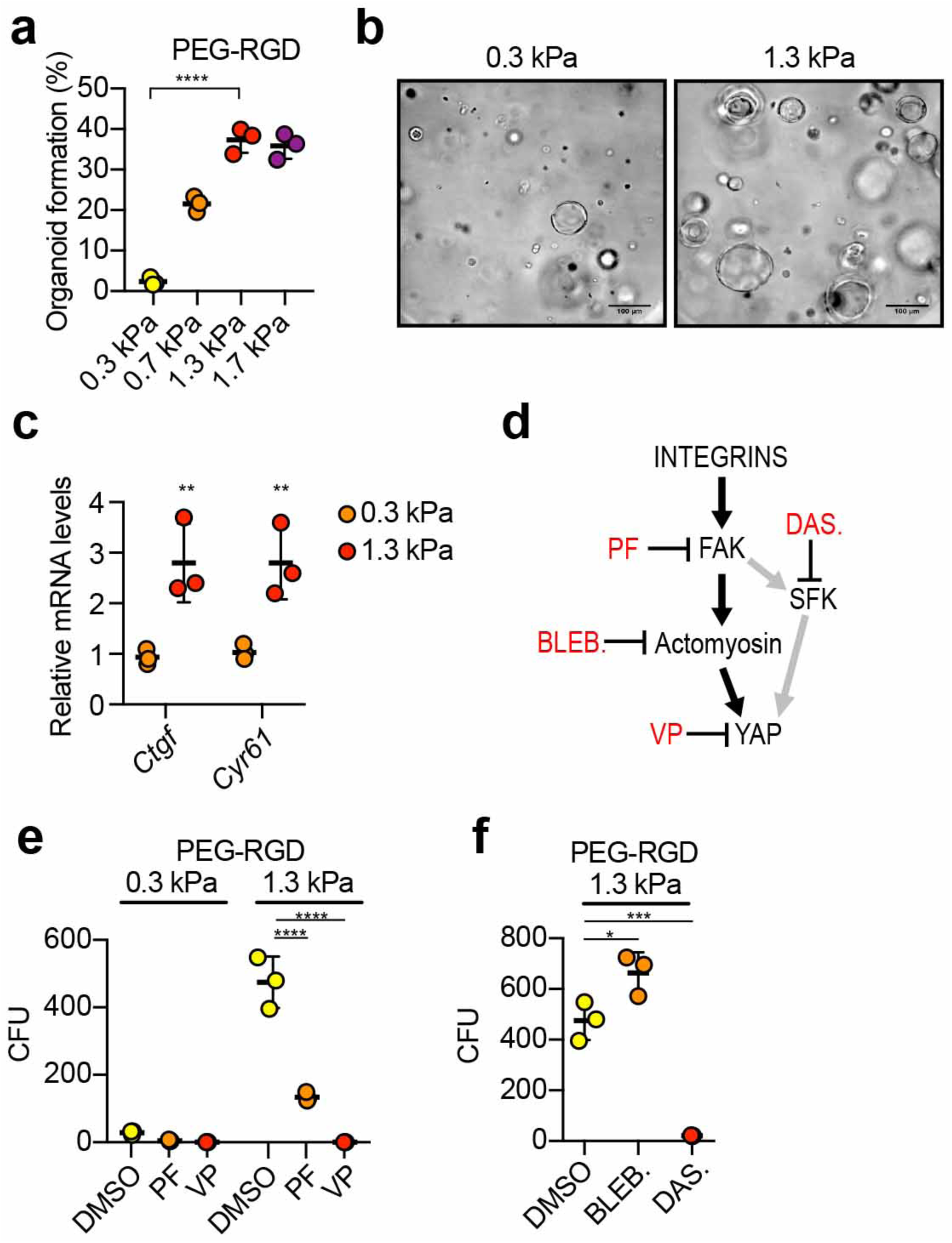
Effect of matrix stiffness on liver organoid formation. **(a)** Effect of matrix stiffness on organoid formation efficiency. **(b)** Representative image of organoids 3 days after embedding in PEG-RGD hydrogels of indicated stiffness. **(c)** Gene expression was analysed by qPCR in liver organoids 6 days after embedding in PEG-RGD hydrogels with indicated stiffness. **(d)** Schematic representation of cellular mechano-signalling pathways. Inhibitors of key elements are depicted in red. **(e)** Effect of indicated inhibitors on organoid formation efficiency in soft (300 Pa) and physiologically-stiff (1.3 kPa) PEG-RGD hydrogels. CFU (colony forming unit). **(f)** Effect of indicated inhibitors on organoid formation efficiency in physiologically-stiff (1.3 kPa) PEG-RGD hydrogels. CFU (colony forming unit). Graphs show individual data points derived from n=3 independent experiments and means ± s.d. *P<0.05, ** P<0.01 ***P<0.001 one way Anova (a, f) or Two way Anova (c, e).

Given the pivotal role of the Hippo pathway nuclear effector, Yes associate protein (YAP), in the transduction of micro-environment mechanical cues downstream of integrins^25^, we next tested whether YAP activation could potentially explain the observed matrix stiffness dependence. We monitored the expression of canonical YAP target genes and YAP subcellular localization in organoids cultured in soft (0.3 kPa) and physiologically-stiff (1.3 kPa) matrices. Interestingly, in stiffer hydrogels, YAP target genes expression and nuclear accumulation were increased compared to soft matrices (Fig. 3c and Supplementary Fig. 3c). To examine whether an activated integrin/YAP signalling axis is functionally required for organoid derivation in physiologically-stiff matrices, we treated organoid cultures with PF-573228, an inhibitor of the integrin effector focal adhesion kinase (FAK), or with the YAP inhibitor Verteporfin^26^ (Fig. 3d). Both treatments prevented the increase in organoid formation induced in stiffer matrices (Fig. 3e and Supplementary Fig. 3d), indicating that the integrin/YAP module is required in coordinating growth of liver progenitors in response to mechanical stimuli.

We then sought to identify the other components of the FAK-YAP cascade that may play a role in modulating the stiffness response in our system. Since remodelling of the actin cytoskeleton has been identified as a key event upstream of YAP activation^27–29^, we assessed its putative involvement in physiological stiffness-induced organoid growth by inhibiting acto-myosin contractility with blebbistatin^30^. Surprisingly, blebbistatin treatment significantly enhanced organoid formation (Fig. 3f), indicating that matrix stiffness promotes organoid growth independently of cytoskeletal dynamics, and that acto-myosin contractility rather interferes with normal liver progenitor expansion. However, tyrosine phosphorylation of YAP by the Scr family of kinases (SFK), an alternative route for integrin-dependent and acto-myosin independent YAP activation^31,32^, was increased by matrix stiffness (Supplementary Fig. 3e). Of interest, treatment with Dasatinib^33^, a FDA-approved SFK inhibitor, fully abolished YAP phosphorylation and organoid growth in physiologically-stiff (1.3 kPa) matrices (Fig. 3f and Supplementary Fig. 3e). These results corroborate the importance of the integrin/SFK/YAP signalling pathway in liver progenitor proliferation in response to differential mechanical inputs.

Liver disease progression is strongly associated with abnormal tissue architecture and mechanotransduction^15,34,35^. Indeed, as a direct effect of aberrant ECM deposition in the fibrotic liver, tissue stiffness increases in time and severely compromises its function^36–38^. In fact, the changes in liver stiffness associated with disease are used for diagnostics based on longitudinal non-invasive monitoring^39^. We reasoned that liver organoids grown in defined hydrogels recapitulating the stiffness of fibrotic liver could serve as physiologically relevant 3D model to investigate how stem cells translate aberrant mechanical inputs into disease-relevant phenotypes. To this aim, we generated fibrosis-mimicking hydrogels with a stiffness of 4 kPa^15,16^. Strikingly, these hydrogels led to a significant impairment of organoid formation (Figs. 4a-b), demonstrating that an abnormal ECM stiffness is sufficient to decrease the liver progenitor proliferative capacity. In this condition liver organoids showed a reduction in the expression of hepatic progenitor markers (Fig. 4c) and upregulation of genes involved in cellular response to hepatic injury (Fig. 4d), indicating impaired stemness potential and concomitant induction of a stress response. Finally, fibrosis-mimicking hydrogels led to an increase in the expression of matrix metalloproteases (Figure 4e), a compensatory phenomenon known to be induced in response to increased ECM stiffness^40–42^. Surprisingly, in these conditions YAP activation was not affected (Fig. 4f), suggesting the existence of other pathways controlling liver progenitor growth in response to increased stiffness. These results suggest that synthetic hydrogels may be a useful tool to assess the contribution of mechanical inputs on liver diseases.

**Figure 4.**
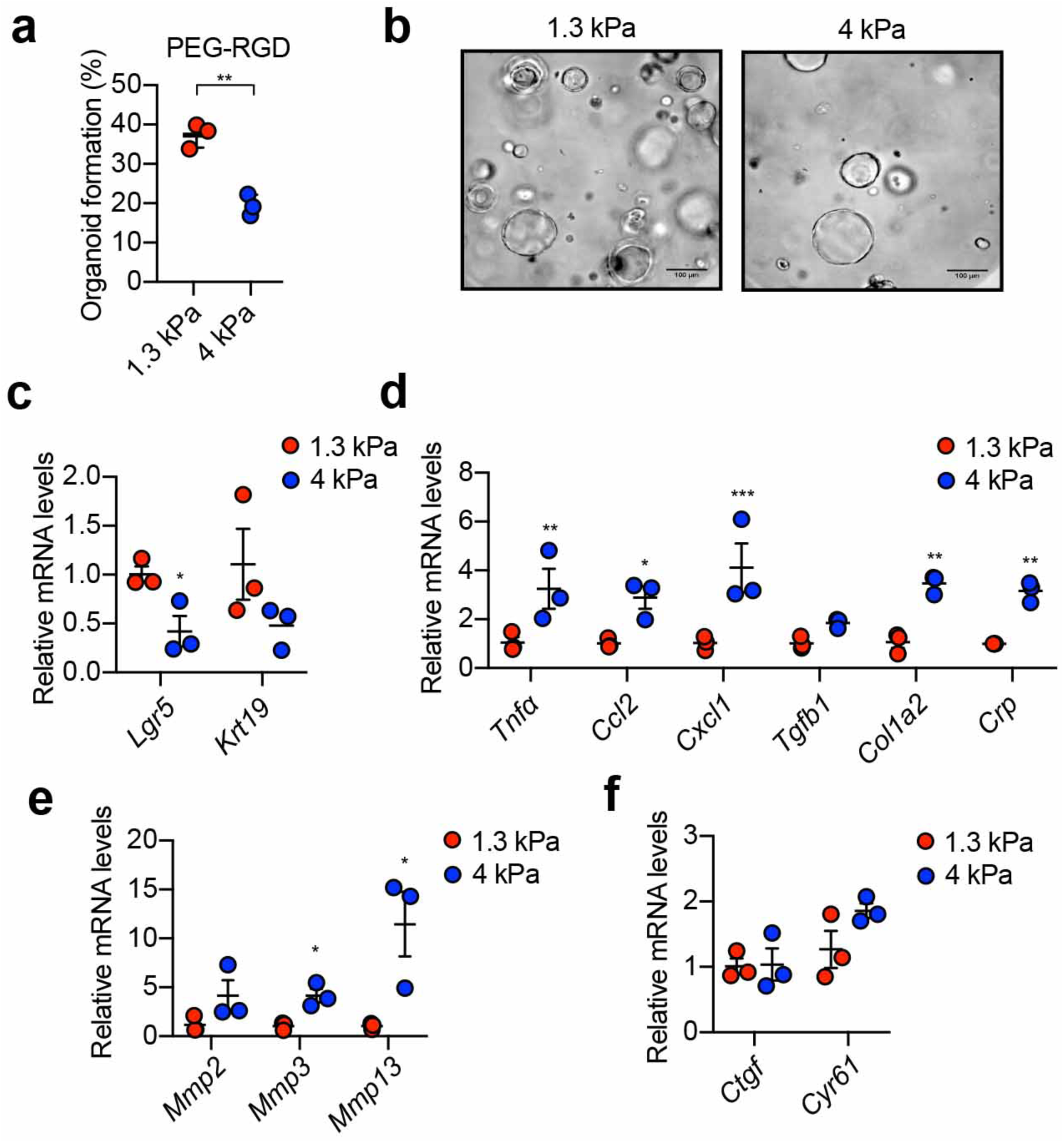
Hydrogels mimicking the stiffness of native fibrotic liver affect the growth of liver organoids and promote a stress response. (**a)** Effect of matrix stiffness on organoid formation efficiency. Scale bars: 100 μm. **(b)** Representative image of organoids 3 days after embedding in PEG-RGD hydrogels of indicated stiffness. (**c-f)** Gene expression was analysed by qPCR in liver organoids 6 days after embedding in PEG-RGD hydrogels with indicated stiffness. Graphs show individual data points derived from n=3 independent experiments and means ± s.d. *P<0.05, ** P<0.01 ***P<0.001 unpaired Student’s two-tailed t-test (a) or two way Anova (c-f).

To test the potential clinical relevance of our findings, we assessed whether the PEG-RGD hydrogels, initially designed for expansion and differentiation of mouse liver organoids, were also suitable for culturing human organoids. We first generated Matrigel-derived organoids from human non-tumorigenic liver needle biopsies^1,43^. Human liver progenitor cells were then embedded in PEG-RGD and cultured in human expansion medium (HEM) (Supplementary Fig. 4a). In these conditions human liver progenitor cells generated organoids that could be expanded over multiple passages (Fig. 5a). Similar to mouse, human organoid generation efficiency in PEG-RGD was comparable to Matrigel (Fig. 5a), and was critically dependent on changes in hydrogel stiffness (Fig. 5b). Moreover, when cultured in HEM, human liver organoids grown in PEG-RGD expressed progenitor cell markers, such as KRT19 (Fig. 5c), but readily differentiated into human hepatocyte-like cells when cultured in human differentiation medium (HDM) (Fig. 5d and Supplementary Fig. 4b).

**Figure 5.**
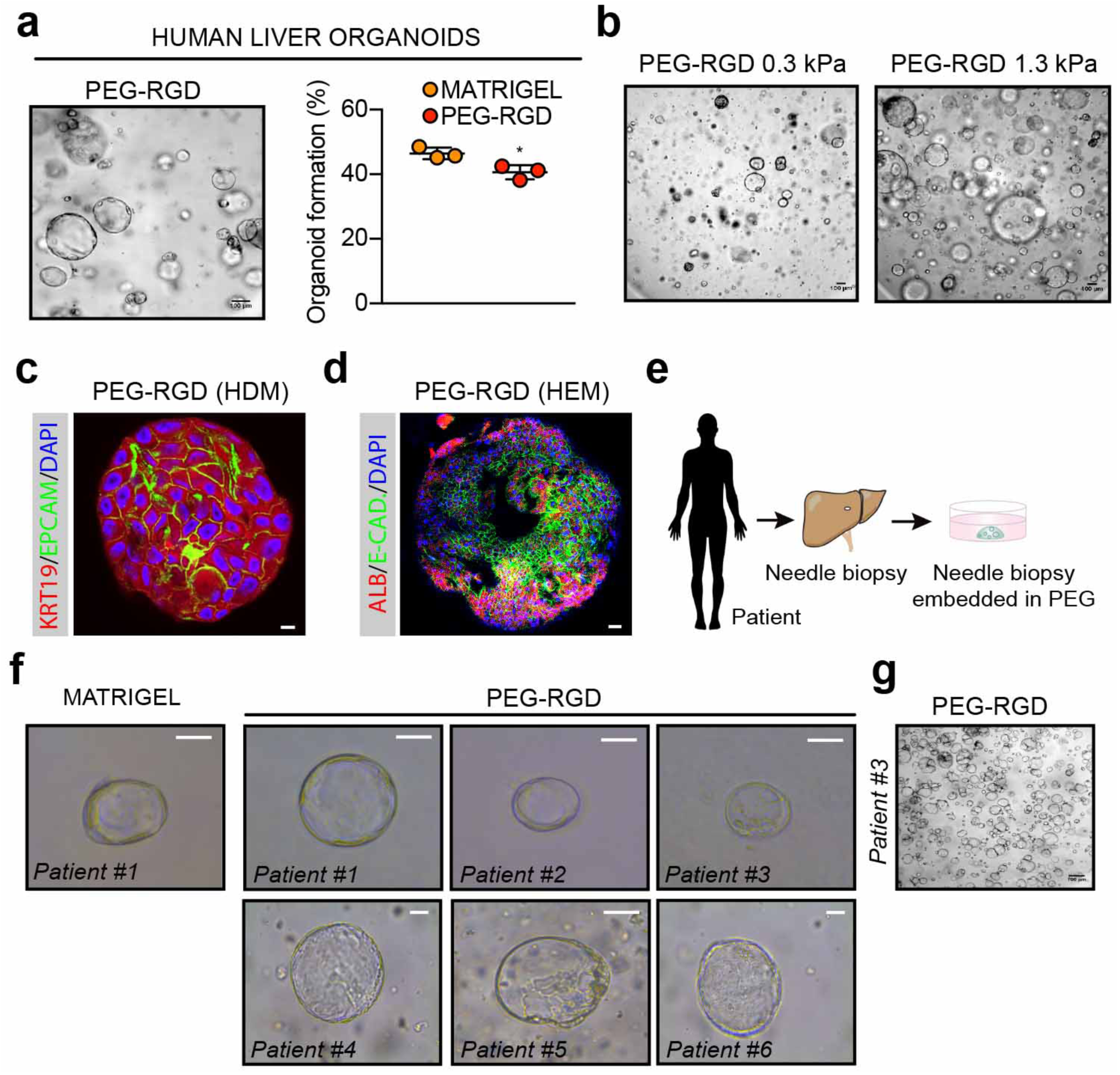
PEG-RGD hydrogels allow for establishment, expansion and differentiation of patient-derived human liver organoids. **(a)** Matrigel-derived human liver progenitor cells were embedded in physiologically-stiff PEG-RGD hydrogels and passaged in human expansion medium (HEM) with an efficiency comparable to Matrigel. **(b)** Effect of matrix stiffness on human liver organoid formation. **(c)** Representative confocal immunofluorescence image of KRT19 and Epcam in liver organoids embedded in physiologically-stiff PEG-RGD hydrogels and cultured in HEM. Scale bar 10 μm. **(d)** Representative confocal immunofluorescence image of Albumin and E-Cadherin in liver organoids cultured in human differentiation medium (HDM), Scale bar 25 μm. **(e, f)** Freshly-isolated liver biopsies from six patients were digested and directly embedded in PEG-RGD hydrogels. Scale bar 50 μm. **(g)** PEG-RGD-derived human liver progenitor cells were embedded in physiologically-stiff hydrogels and passaged in human expansion medium (HEM). Scale bar 100 μm. *P<0.05, unpaired Student’s two-tailed t-test.

Finally, in order to generate clinically relevant human liver organoids, we tested the possibility of establishing organoids from human patients without the interference of any animal-derived matrices. To this aim, freshly isolated liver biopsies from six patients were digested and directly embedded in PEG-RGD hydrogels (Fig. 5e). Strikingly, after 8 days of culture, organoid formation was scored from progenitor cells of all patients (Fig. 5f), and could be passaged (Fig. 5g). Altogether, these data provide proof-of-concept that human liver progenitor cells can be expanded and maintained *in vitro* within synthetic matrices.

## Discussion

We report here the establishment of a fully defined 3D culture system for mouse and human hepatic progenitors and organoids. We demonstrated that liver organoids can be expanded and maintained in such minimal environments with an efficiency that is comparable to Matrigel, but without its main disadvantages of structural instability, batch-to-batch variability and clinical incompatibility. By tuning the stiffness of the synthetic networks to match the physiological levels of the liver, we optimized the efficiency of liver organoid derivation, and identified integrin-SFK-YAP as a mechano-sensitive axis that is required for liver organoid growth. Interestingly, in contrast to intestinal organoids^7^, we found that liver progenitor cells transduce mechanical signals in an acto-myosin independent manner and instead require activation of the tyrosine kinase Src to support epithelial tissue formation. Moreover, we used PEG hydrogels to accurately model the aberrant mechanical properties of the fibrotic liver, providing evidence that aberrant liver stiffness negatively impacts liver progenitor proliferation. This experimental set-up provides a standardized framework to study hepatic progenitor cells in a defined mechanical environment, which may further our understanding of the underappreciated role of mechanical cues in modulating the molecular properties and signatures of this cell population in healthy and fibrotic liver. Finally, our data showing that clinically relevant human stem/progenitor cells can be grown *in vitro* without any requirement of animal-derived matrices, may open exciting perspectives for the establishment of protocols for liver organoid-based clinical applications.

## Acknowledgments

We thank Andréane Fouassier, Sabrina Bichet, Thibaud Clerc, Laure Vogeleisen-Delpech, Fabiana Fraga, the Phenotyping Unit (UDP) and the Histology core facility (HCF) of EPFL for technical assistance. The work of K.S. was funded by the Swiss National Science Foundation (SNSF 31003A_125487), Sinergia CRSII3_160798/1 and the Ecole Polytechnique Fédérale de Lausanne (EPFL). The work of M.P.L. in the area of organoid biology and technology was supported by the Swiss National Science Foundation (grant #310030_179447), the European Union’s Horizon 2020 research and innovation programme (INTENS 668294), the Personalized Health and Related Technologies Initiative from the ETH Board, the Vienna Science and Technology Fund and École Polytechnique Fédérale de Lausanne (EPFL). G.S. was funded by a postdoctoral FEBS long-term fellowship.

## Author contributions

G.S., S.R., M.L. and K.S. conceived the project and wrote the manuscript. G.S., S.R. and E.Y. planned and performed experiments and analysed data. S.N. and M.H. provided human biopsy samples and critically revised the manuscript.

## Methods

### Data reporting

The experiments were not randomized and the investigators were not blinded to allocation during experiments and outcome assessment. No statistical methods were used to determine sample size. The experiments were repeated at least 3 times or with at least 3 different donors to control biological variations.

### Animals and ethical approval

Liver tissues were harvested from 8-12 weeks old euthanized C57BL/6J male mice. All the animal experiments were authorized by the Veterinary Office of the Canton of Vaud, Switzerland under the license authorization no. 3263.

### Enzymatically crosslinked hydrogel precursor synthesis

Hydrogel precursors were synthesized as previously reported^44^. Briefly, vinylsulfone functionalized 8-arm PEG (PEG-VS) was purchased from NOF. The transglutaminase (TG) factor XIII (FXIIIa) substrate peptides Ac-FKGG*GPQGIWGQ*-ERCG-NH2 with matrix metalloproteinases (MMPs) sensitive sequence (in italics), Ac-FKGG-GDQGIAGF-ERCG-NH2, H-NQEQVSPLERCGNH2 and the RGD-presenting adhesion peptide H-NQEQVSPLRGDSPG-NH2 were purchased from GL Biochem. FXIIIa substrate peptides and 8-arm PEG-VS were dissolved in triethanolamine (0.3 M, pH 8.0) and mixed at 1.2 stoichiometric excess (peptide-to-VS group), and allowed to react for 2 h under inert atmosphere. The reaction solution was dialysed (Snake Skin, MWCO 10K, PIERCE) against ultrapure water for 3 days at 4 °C, after which the products were lyophilized and dissolved in ultra-pure water to make 13.33% w/v stock solutions.

### Formation and dissociation of PEG hydrogels

PEG precursor solutions were mixed in stoichiometrically balanced ratios to form hydrogel networks of a desired final PEG content. Addition of thrombin-activated FXIIIa (10 U ml−1; Galexis) triggered the hydrogel formation in the presence of Tris-buffered saline (TBS; 50 mM, pH 7.6) and 50 mM CaCl2. The spare reaction volume was used for the incorporation of dissociated liver stem cells, fragments of liver bile ducts, and ECM components: RGD-presenting adhesion peptide, fibronectin (0.5 mg ml−1; R&D systems), laminin-111 (0.2 mg ml−1; Invitrogen), collagen-IV (0.2 mg ml−1; BD Bioscience). Gels cast on PDMS-coated 24 well-plate were allowed to crosslink by incubation at 37°C for 10 min. To release the grown colonies for further processing, gels were detached from the bottom of the plates using a tip of a metal spatula and transferred to 15-ml Falcon tube containing 1 ml of Dispase (1 mg/ml, Thermo Fisher Scientific). After 10 minutes enzymatic digestion, the reaction was quenched using 10% FBS containing 1 mM EDTA, washed with cold basal medium and centrifuged for 3 min at 1000 rpm.

### Mechanical characterization of PEG hydrogels

Elastic modulus (G’) of hydrogels was measured by performing small-strain oscillatory shear measurements on a Bohlin CVO 120 rheometer with plate-plate geometry. Briefly, 1-1.4 mm thick hydrogel discs were prepared and allowed to swell in water for 24 hrs. The mechanical response of the hydrogels sandwiched between the parallel plates of the rheometer was recorded by performing frequency sweep (0.1-10 Hz) measurements in a constant strain (0.05) mode at 25 °C.

### Quantification of liver organoid formation efficiency

Phase contrast z-stacks images were collected through the entire thickness of the PEG gels (every 15 µm) at 4 different locations within the gels (Nikon Eclipse Ti). The Cell Counter plugin in ImageJ (NIH) was used to quantify the percentage of single cells that formed colonies after 3 days of culture in expansion medium.

### Culture of mouse and human liver organoids from biliary duct fragments and single cells

Mouse liver organoids were established from biliary duct fragments as previously described with some modifications^2^. Briefly, liver tissues were digested in digestion solution (Collagenase type XI 0.012%, dispase 0.012%, FBS 1% in DMEM medium) for 2 hours. When digestion was complete, bile ducts were pelleted by mild centrifugation (200 rpm for 5 min) and washed with PBS. Isolated ducts were then resuspended either in Matrigel (BD Bioscience) or PEG precursor solution and cast in 10 µl droplets in the center of the wells in a 48-well plate. After the gels were formed, 250 µl of isolation medium was added to each well. Isolation medium was composed of AdDMEM/F12 (Invitrogen) supplemented with B27 and N2 (both GIBCO), 1.25 µM N-acetylcysteine (Sigma-Aldrich), 10 nM gastrin (Sigma-Aldrich) and the following growth factors: 50 ng ml^−1^ EGF (Peprotech), 1 µg ml^−1^ Rspo1 (produced in-house), 100 ng ml^−1^ Fgf10 (Peprotech), 10 mM nicotinamide (Sigma-Aldrich), 50 ng ml^−1^ HGF (Peprotech), Noggin (100 ng ml^−1^ produced in-house), Wnt 3a (1µg ml^−1^, Peprotech) and Y-27632 (10 µM, Sigma). After the first 4 days, isolation medium was changed with expansion medium (EM) which consists of isolation medium without Noggin, Wnt and Y-27632. One week after seeding, organoids were removed from the Matrigel or PEG hydrogel, dissociated into single cells using TrypLE express (Gibco), and transferred to fresh Matrigel or PEG hydrogels. Passaging was performed in 1:3 split ratio once per week. Plasmids for Rspo1 and Nog production were a kind gift from Joerg Huelsken. Liver progenitor cells were treated with the following compounds for YAP-inhibition experiments: Verteporfin (10 µM, Sigma), Dasatinib (10 µM, Selleckchem), Blebbistatin (10 µM StemCell Technologies), PF 573228 (10 µM Tocris).

### Human liver biopsies and generation of human organoids

Human tissues were obtained from patients undergoing diagnostic liver biopsy at the University Hospital Basel. Written informed consent was obtained from all patients. The study was approved by the ethics committee of the northwestern part of Switzerland (Protocol Number EKNZ 2014-099). Ultrasound (US)-guided needle biopsies were obtained with a coaxial liver biopsy technique as described previously^43^. One biopsy cylinder was fixed in formalin and paraffin-embedded for histopathological diagnosis. Additional cylinders were collected in advanced DMEM/F-12 (GIBCO) for organoid generation. Patient clinical information are shown in Supplementary Table 2. Human liver organoids were generated as previously described with some modifications^43^. Briefly, biopsies were placed in advanced DMEM/F-12 (GIBCO) and transported to the laboratory on ice. Liver samples were then digested to small-cell clusters in basal medium containing 2.5 mg/mL collagenase IV (Sigma) and 0.1 mg/mL DNase (Sigma) at 37°C. Cell clusters were embedded in Matrigel or PEG gels, cast and after the gels were formed, human isolation medium (HIM) was added. HIM is composed of advanced DMEM/F-12 (GIBCO) supplemented with B-27 (GIBCO), N-2 (GIBCO), 10 mM nicotinamide (Sigma), 1.25 mM N-acetyl-L-cysteine (Sigma), 10 nM [Leu15]-gastrin (Sigma), 10 µM forskolin (Tocris), 5 µM A83-01 (Tocris), 50 ng/mL EGF (PeproTech), 100 ng/mL FGF10 (PeproTech), 25 ng/ml HGF (PeproTech), 1 µg ml^−1^ Rspo1 (produced in-house), Wnt3a (1µg ml^−1^, Peprotech), Y-27632 (10 µM, Sigma) and Blebbistatin (10 µM StemCell Technologies). After the first 4 days, isolation medium was changed with human expansion medium (HEM) which consists of HIM without Noggin, Wnt and Y-27632.

### Mouse hepatocyte differentiation

Single cells were seeded and kept for 6 days in EM. Then the medium was changed to differentiation medium (DM) which no longer contains Rspo1, HGF and nicotinamide and instead contains A8301 (50 nM, Tocris Bioscience) and DAPT (10 nM, Sigma-Aldrich). Cells were maintained in DM for 12 days. During the last 3 days DM was also supplemented with dexamethasone (Sigma, 3 µM). Medium was changed every two days.

### Human hepatocyte differentiation

Single cells were seeded and kept 6 days in HEM. Then the medium was changed to differentiation medium (HDM) which no longer contains Rspo1, HGF and nicotinamide and instead contains A8301 (50 nM, Tocris Bioscience) and DAPT (10 nM, Sigma-Aldrich), BMP7 (25ng/ml Peprotech) and human Fgf19 (100 ng/ml, R&D). Cells were maintained in DM for 10 days. During the last 3 days DM was also supplemented with dexamethasone (3 µM). Medium was changed every two days.

### Immunohistochemical analysis of human organoids

Human organoids were fixed in 10% neutral buffered formalin, washed with PBS, dehydrated, and embedded in paraffin. 5 µm thick sections were made from paraffin-embedded samples and sections were stained with H&E and PAS.

### Immunofluorescence analysis

Liver organoids were extracted from Matrigel (with Cell Recovery Solution, Corning) or PEG gels (with 1 mg ml− 1 Dispase (Gibco) for 10 min at 37 °C) and fixed with 4% paraformaldehyde (PFA) in PBS (20 min, room temperature). Organoids in suspension were centrifuged (1000 r.p.m., 5 min) to remove the PFA, washed with ultra-pure water and pelleted. The organoids were then spread on glass slides and allowed to attach by drying. Attached organoids were rehydrated with PBS and permeabilized with 0.2% Triton X-100 in PBS (1 h, room temperature) and blocked (1% BSA in PBS) for 1 h. Samples were then incubated overnight with phalloidin-Alexa 488 (Invitrogen) and primary antibodies against Epcam (1:50, eBioscience, G8.8), Krt19 (1:100, ab15463), E-cadherin (1:100, Cell Signaling, 24E10), Sox9 (1:50, Millipore, AB5535), Albumin (1:50, R&D systems, MAB1455), Hnf4α (1:50, Santa Cruz, C19), ZO-1 (1:50, Invitrogen, 61-7300); YAP (1:50, Cell Signaling, 4912S). Samples were washed with PBS and incubated for 3 h with secondary antibodies Alexa 488 donkey-α-rabbit, Alexa 568 donkey-α-mouse, Alexa 647 donkey-α-goat (1:1000 in blocking solution; Invitrogen). Following extensive washing, stained organoids were imaged by confocal (Zeiss LSM 710) mode. Dapi was used to stain nuclei.

### Western blotting

Samples were lysed in lysis buffer (50 mM Tris (pH 7.4), 150 mM KCl, 1 mM EDTA, 1% NP-40, 5 mM NAM, 1 mM sodium butyrate, protease and phosphatase inhibitors). Proteins were separated by SDS–PAGE and transferred onto nitrocellulose or polyvinylidene difluoride membranes. Blocking (30 min) and antibody incubations (overnight) were performed in 5% BSA in TBST. YAP1 (sc101199, 1:1000), phospho-YAP1^Y357^(ab62751, 1:1000), actin (sc47778, 1:1000).

### Quantitative real-time qPCR for mRNA quantification

Liver organoids were extracted from Matrigel or PEG gels as previously described^7^. RNA was extracted from organoids using the RNAqueous total RNA isolation kit (Thermo Fisher) following manufacturer’s instructions. RNA was transcribed to complementary DNA using QuantiTect Reverse Transcription Kit (Qiagen) following manufacturer’s instructions. Expression of selected genes was analysed using the LightCycler 480 System (Roche) and SYBR Green chemistry. Quantitative polymerase chain reaction (PCR) results were presented relative to the mean of Gapdh (DDCt method). Quantitative polymerase chain reaction (PCR) results shown as heat map instead represent the DCt values. Primers for qPCR are listed in Supplementary Table 1.

### Proliferation assay

Cell proliferation was assessed by EdU assay (Click-iT EdU Alexa Fluor 647) following manufacturer’s instructions.

### Functional analysis

LDL uptake was detected with DiI-Ac-LDL (Biomedical Technologies). Mouse albumin secretion was detected with ELISA kit (Abcam, ab108792). Urea secretion was assessed with QuantiChrom™ Urea Assay Kit (BioAssay Systems). All experiments were followed according to the manufacturers’ instructions.

### Statistical analysis and sample information

Statistically significant differences between the means of two groups were assessed by t-test as specified in the legends. All statistical analyses were performed in the GraphPad Prism 7.0 software. A P value of 0.05 or less was considered statistically significant.

### Data availability

All summary or representative data generated and supporting the findings of this study are available within the paper. Raw data that support the findings of this study are available upon reasonable request.

---

**Supplementary Figure 1.**
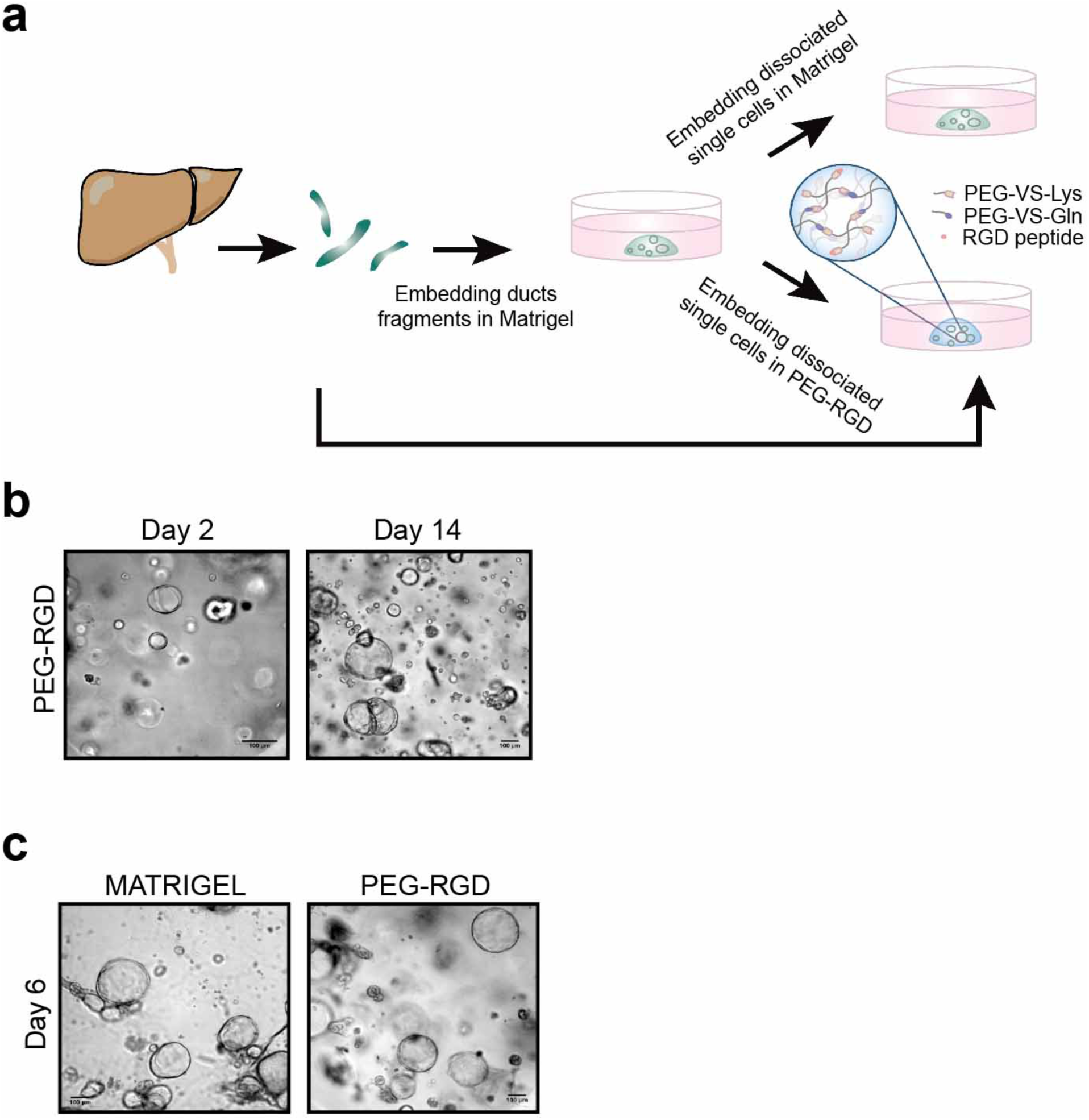
**(a)** Schematic of the protocol for generating mouse liver organoid cell lines in Matrigel or, directly, in PEG-RGD hydrogels. **(b)** Representative picture of human liver organoids embedded in PEG-RGD for the indicated times. **(c)** PEG-RGD hydrogel provides a stable matrix for long term culture of hepatic organoids, while Matrigel already softened after 6 days of culture. Scale bars: 100 μm.

**Supplementary Figure 2.**
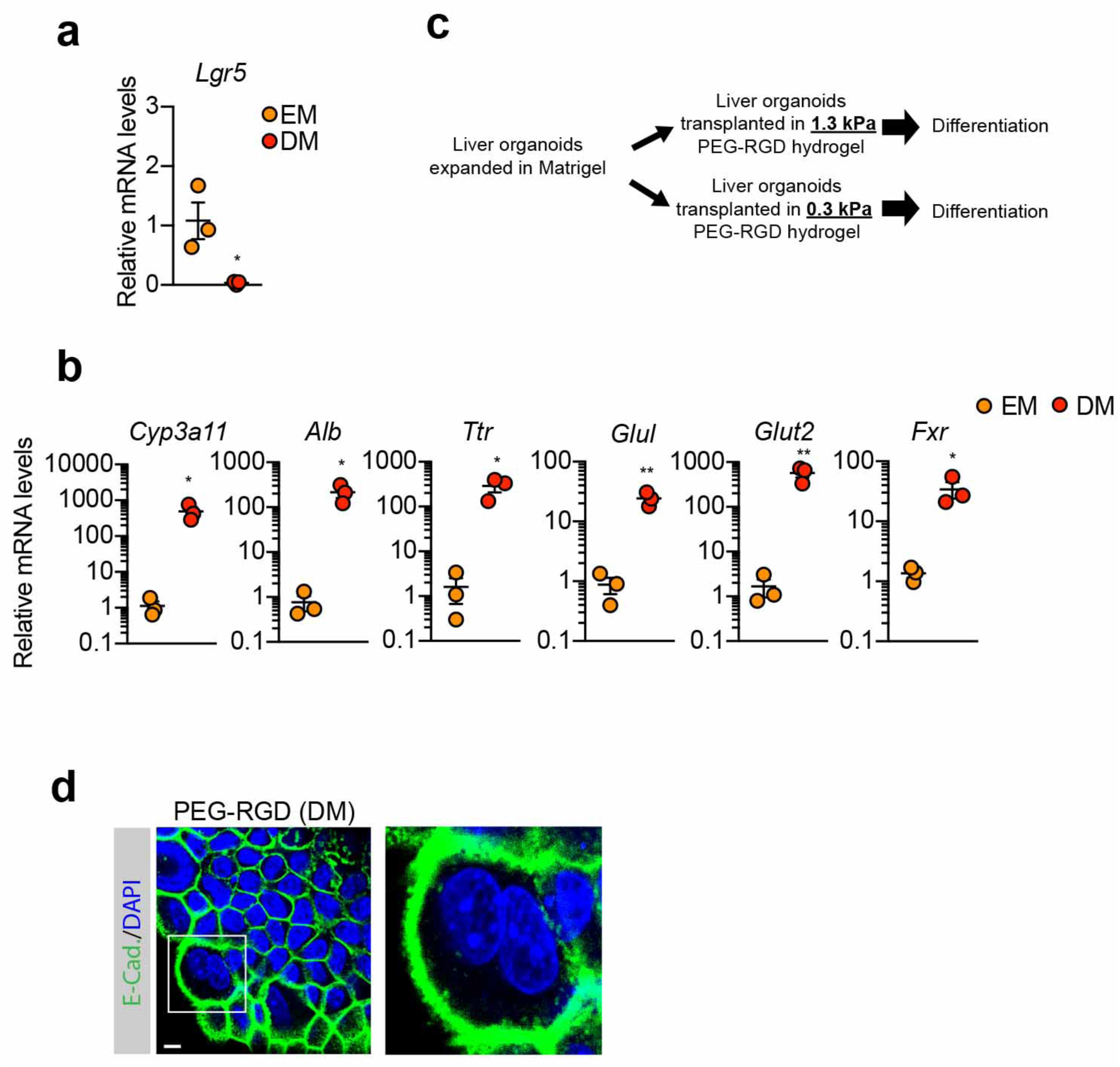
**(a)** Lgr5 mRNA levels were analysed by qPCR in liver organoids maintained in differentiation medium (DM). Graph show individual data points derived from n=3 independent experiments and means ± SEM. **(b)** Gene expression was analysed by qPCR in liver organoids maintained in expansion medium (EM) or differentiation medium (DM) in Matrigel. Graphs show individual data points derived from n=3 independent experiments and means ± SEM. *P<0.05, ** P<0.01 paired Student’s two-tailed t-test. **(c)** Liver organoids grown in physiologically-stiff hydrogels were transplanted in soft hydrogels immediately before inducing the differentiation. **(d)** Liver organoids maintained in DM in PEG-RGD hydrogels. E-cadherin was used to visualize cell borders. The square indicates a bi-nucleated cell. Scale bar 10 μm. Graphs show individual data points derived from n=3 independent experiments and means ± SEM. *P<0.05, unpaired Student’s two-tailed t-test.

**Supplementary Figure 3.**
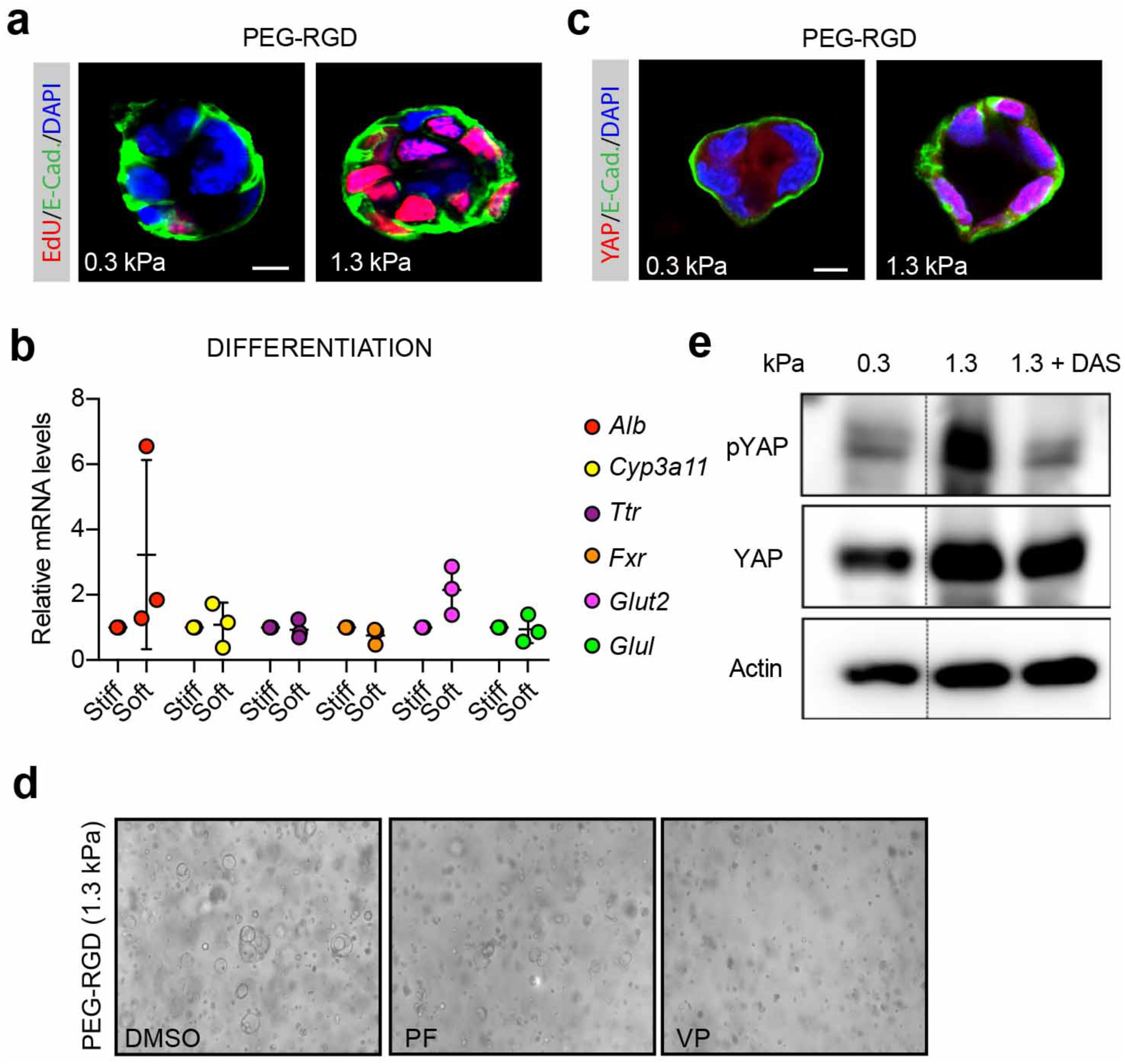
**(a)** Representative images of EdU staining of liver organoids, 3 days after embedding in PEG-RGD hydrogels with indicated stiffness. **(b)** Gene expression was analysed by qPCR in liver organoids embedded in soft (0.3 kPa) or physiologically-stiff (1.3 kPa) PEG-RGD hydrogels and maintained in DM. Data are shown in relative to the expression in physiologically-stiff hydrogels. **(c)** Representative image of YAP subcellular localization in liver organoids 1 day after embedding in soft (0.3 kPa) or physiologically-stiff (1.3 kPa) PEG-RGD hydrogels. **(d)** Representative images relative to Fig. 3e. **(e)** Western blot showing YAP phosphorylation in mouse liver progenitor cells 6 hours after embedding in soft (0.3 kPa) and physiologically-stiff (1.3 kPa) PEG-RGD hydrogels.

**Supplementary Figure 4.**
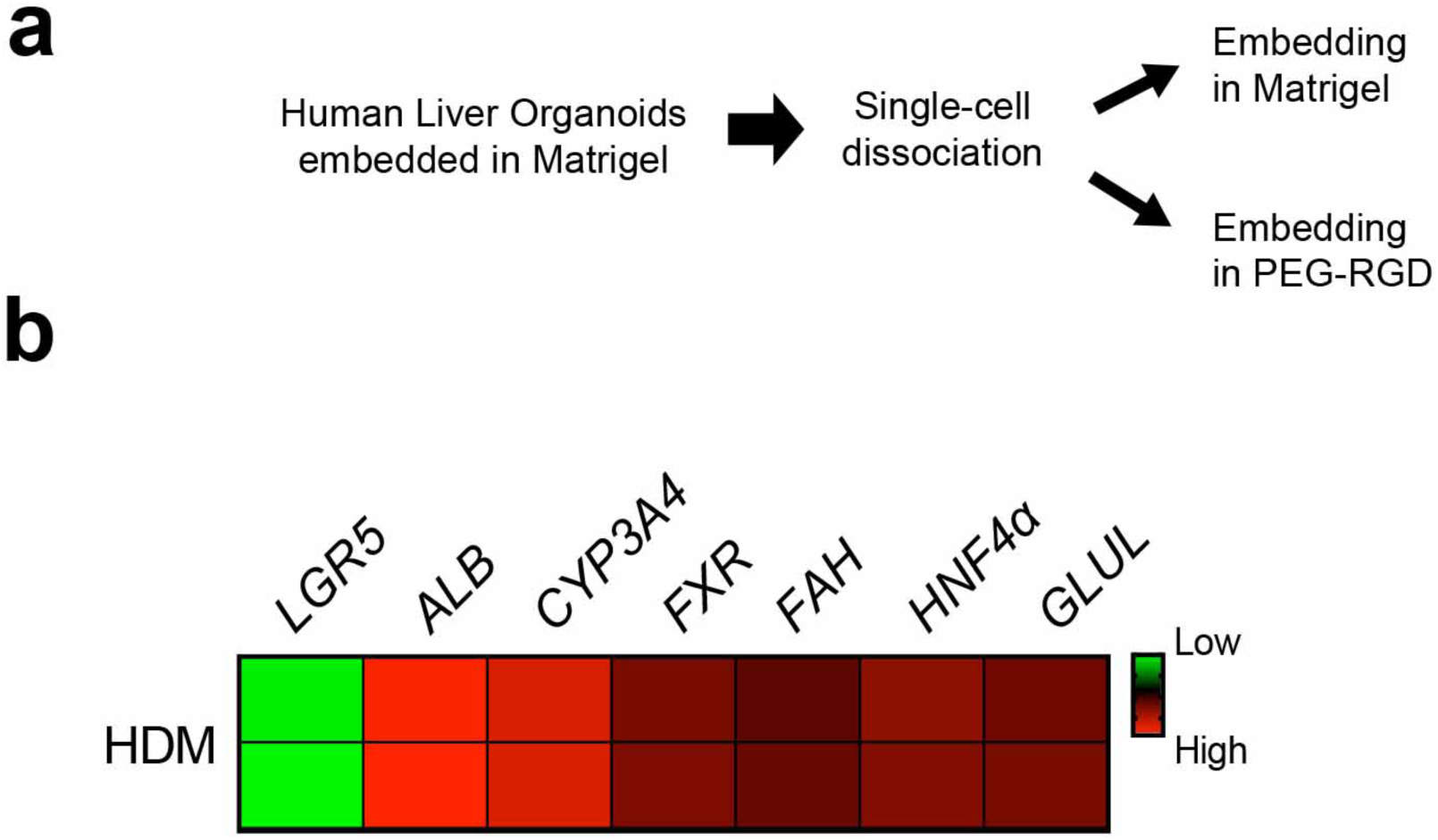
**(a)** Schematic representation of the protocol. **(b)** Gene expression was analysed by qPCR in human liver organoids embedded in PEG-RGD hydrogels and maintained in human differentiation medium. The heat-map shows results from two biological replicates.

**Supplementary Table 1.**
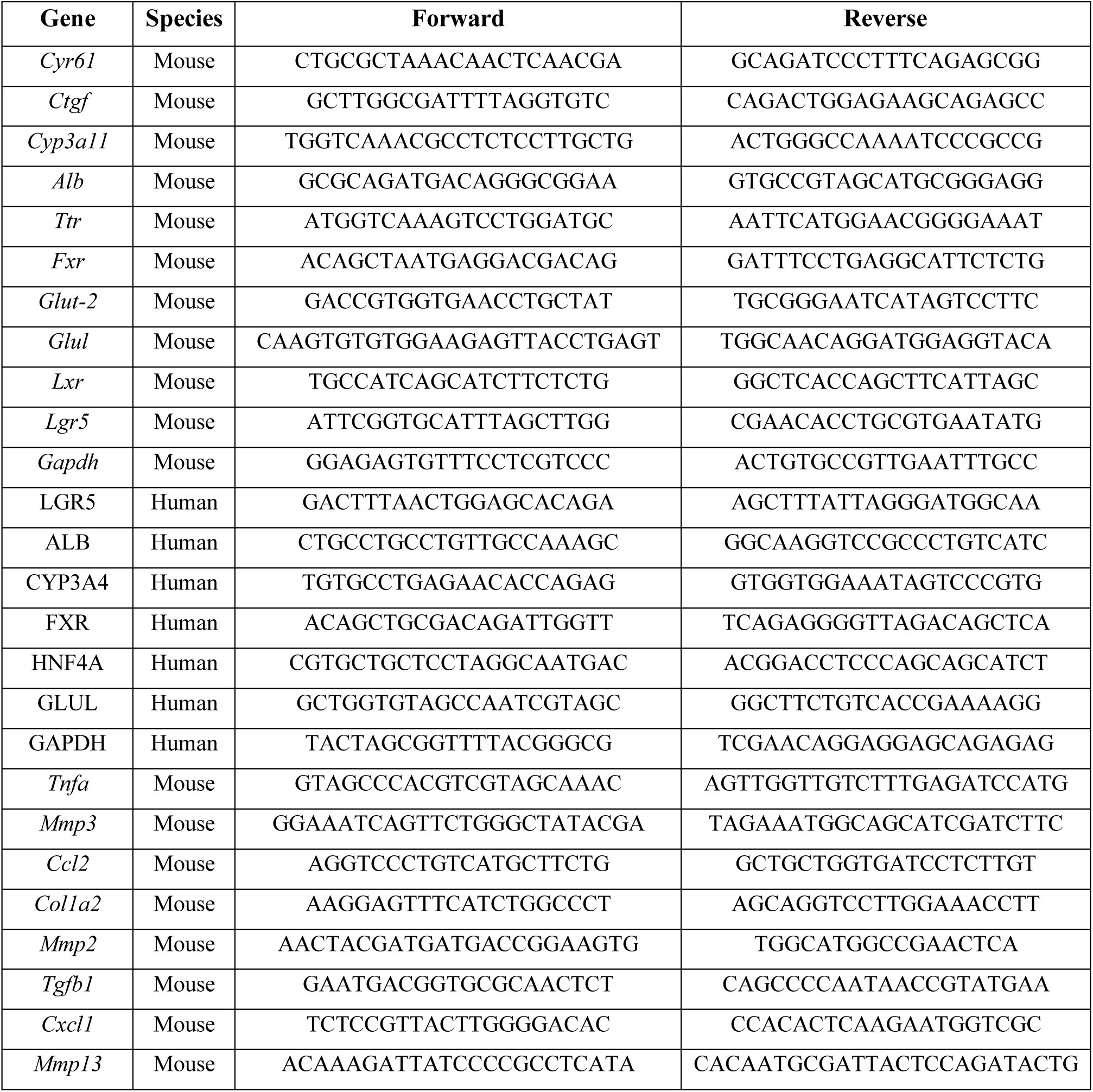
Primers used for qPCR analysis.

**Supplementary Table 2.** Patient clinical information.

| **Sex** | **Age** | **Diagnosis** | **Cirrhosis** |
| --- | --- | --- | --- |
| Male | 58 | ASH, alcoholic steatohepatitis | YES |
| Female | 36 | AFLD, alcoholic fatty liver disease | NO |
| Male | 43 | NAFLD, nonalcoholic fatty liver disease | NO |
| Female | 45 | NASH, nonalcoholic steatohepatitis | NO |
| Male | 35 | NASH, nonalcoholic steatohepatitis | NO |
| Female | 73 | Normal | NO |
| Male | 64 | ASH/NASH | NO |

